# Robust methods for detecting convergent shifts in evolutionary rates

**DOI:** 10.1101/457309

**Authors:** Raghavendran Partha, Amanda Kowalczyk, Nathan Clark, Maria Chikina

## Abstract

Identifying genomic elements underlying phenotypic adaptations is an important problem in evolutionary biology. Comparative analyses learning from convergent evolution of traits are gaining momentum in accurately detecting such elements. We previously developed a method for predicting phenotypic associations of genetic elements by contrasting patterns of sequence evolution in species showing a phenotype with those that do not. Using this method, we successfully demonstrated convergent evolutionary rate shifts in genetic elements associated with two phenotypic adaptations, namely the independent subterranean and marine transitions of terrestrial mammalian lineages. Our method calculates gene-specific rates of evolution on branches of phylogenetic trees using linear regression. These rates represent the extent of sequence divergence on a branch after removing the expected divergence on the branch due to background factors. The rates calculated using this regression analysis exhibit an important statistical limitation, namely heteroscedasticity. We observe that the rates on branches that are longer on average show higher variance, and describe how this problem adversely affects the confidence with which we can make inferences about rate shifts. Using a combination of data transformation and weighted regression, we have developed an updated method that corrects this heteroscedasticity in the rates. We additionally illustrate the improved performance offered by the updated method at robust detection of convergent rate shifts in phylogenetic trees of protein-coding genes across mammals, as well as using simulated tree datasets. Overall, we present an important extension to our evolutionary-rates-based method that performs more robustly and consistently at detecting convergent shifts in evolutionary rates.

## Introduction

Understanding the relationship between phenotype and genotype is one of the fundamental problems of biology. A mechanistic characterization of this relationship hinges on our ability to define how specific genetic elements contribute to biological processes at the molecular, cellular, and organismal level. Advanced high-throughput sequencing technologies have enabled the development of experimental approaches that have discovered a wealth of genetic elements with putative regulatory roles across tissues (ENCODE Project Consortium, 2012; Andersson *et al.*, 2014; Romanoski *et al.*, 2015). However, identifying the precise biological functions of these discovered elements remains a challenge. Even beyond non-coding elements, the precise biological roles of many protein-coding genes is still poorly understood and many genes with statistical disease associations still lack a mechanistic explanation (Pennacchio *et al.*, 2013; Radivojac *et al.*, 2013; Sa, Sánchez and Huarte, 2013; Shlyueva, Stampfel and Stark, 2014). While experimental validation for functional annotation remains challenging, there is considerable interest in developing new tools that can use existing data resources to further elucidate the function of genetic elements. These approaches have the potential to improve the diagnosis of disease susceptibility as well as the development of therapeutic interventions (Manolio *et al.*, 2009; Esteller, 2011).

Computational approaches learning from patterns of convergent phenotypic evolution across species provide a complementary approach to predict genotype-phenotype associations. The natural world is rife with examples of phenotypic convergence ranging from the independent evolution of flight in birds and mammals to diving in species that transitioned from a terrestrial to marine habitat to loss of complex phenotypes such as eyesight in animals colonizing the subterranean niche. Genome-scale studies aimed at identifying the genetic basis of phenotypic convergence take advantage of the growing availability of whole genome sequences for species across several orders, alongside the development of comparative methods to predict orthologous sequences (Eisen, 1998; Pellegrini *et al.*, 1999; Li *et al.*, 2014). An oft considered approach by such studies is to identify convergence at the molecular level, including substitutions at specific nucleotide or amino acid sites (Zhang and Kumar, 1997; Parker *et al.*, 2013; Stern, 2013; Foote *et al.*, 2015; Thomas and Hahn, 2015; Zou and Zhang, 2015). A complementary strategy to investigate the genetic basis of convergence is to search for convergent changes at the level of larger functional regions rather than specific nucleotide or amino acid sites. Sets of genes associated with a phenotype can respond to convergent changes in the selective pressure on the phenotype through non-identical changes in the same gene, and as such, sites-based methods can fail to detect them. These limitations have encouraged researchers to search for convergent shifts in evolutionary rates of individual protein-coding genes and more recently conserved non-coding elements (Lartillot and Poujol, 2011; Hiller *et al.*, 2012; Chikina, Robinson and Clark, 2016; Marcovitz, Jia and Bejerano, 2016; Prudent *et al.*, 2016). An increased selective constraint can manifest as a slower evolutionary rate whereas faster evolutionary rates can result from a release of constraint or from adaptation. Thus phenotypic associations for genetic elements can be predicted from correlated changes in their evolutionary rates on phylogenetic branches corresponding to the phenotypic change. Example approaches based on evolutionary rates include the Forward/Reverse Genomics methods that have identified protein-coding and non-coding genetic elements showing convergent regression in subterranean mammals, as well as limb-regulatory elements lost in snake lineages (Hiller *et al.*, 2012; Marcovitz, Jia and Bejerano, 2016; Prudent *et al.*, 2016; Roscito et al., 2017).

We previously developed an evolutionary-rates-based method to identify genetic elements showing convergent shifts in evolutionary rates associated with two distinct phenotypic transitions (Chikina, Robinson and Clark, 2016; Partha *et al.*, 2017). Our method calculates gene-specific evolutionary rates using a linear model, and gene-trait associations are inferred using correlations of these rates with the phenotype of interest. A genome-wide scan for protein-coding genes associated with the transition to the marine environment identified hundreds of genes that showed accelerated evolutionary rates on three marine mammal lineages (Chikina, Robinson and Clark, 2016). These accelerated genes were significantly enriched for functional roles in pathways important for the marine adaptation including muscle physiology, sensory systems and lipid metabolism. More recently, using our methods we detected an excess of vision-specific genes as well as enhancers that showed convergent rate acceleration on the branches corresponding to four subterranean mammals (Partha *et al.*, 2017). Such genome-scale efforts, both from our group and others, searching for genetic elements responding to convergent changes in the selective pressures in their environment, are gaining momentum in accurately describing precise genotype-phenotype associations.

Our current evolutionary-rates method has an important statistical limitation, namely strong mean-variance trends in the computed evolutionary rates. The distributions of branch lengths of gene trees in phylogenetic datasets are influenced by the choice of species, divergence from the most recent common ancestor, and species-specific properties such as generation time in addition to gene-specific constraints on the sequence evolution. These factors cause large differences in the average lengths as well as the variance of the branch lengths across the branches studied. In this paper, we illustrate how this limitation can adversely impact the confidence with which we infer phenotypic associations for genetic elements, in particular making them sensitive to certain factors in phylogenomic analyses including choice of taxonomic groups and average rates of sequence divergence on phylogenetic branches showing the convergent phenotype. We demonstrate how introducing long branches in phylogenetic trees via the inclusion of distantly related species impacts the reliable estimation of evolutionary rates using gene trees across mammals, as well using a first-of-its-kind model for simulating gene trees. We present key improvements to our methods that address these limitations and overcome them. The next section New Approaches presents a detailed walk-through of our current approach to calculate relative evolutionary rates, the illustration of mean-variance trends (heteroscedasticity) in these rates, and our methodological updates that correct for the problem of heteroscedasticity in the rates. We subsequently demonstrate the improved reliability in relative rate calculations using our updated method, and more importantly in the robust detection of convergent rate shifts across a range of evolutionary scenarios in real and simulated phylogenetic datasets.

## New Approaches

### Current relative-evolutionary-rates methods for predicting phenotypic associations of genetic elements

Our method infers genetic elements associated with a convergent phenotype of interest based on correlations between that phenotype and the rates of evolution of genetic elements. As input, the phenotype is encoded as a binary trait on a phylogenetic tree, and the evolution of each genetic element is similarly described by phylogenetic trees with the same fixed topology. Figure 1 provides an illustration of our method capturing the convergent acceleration of the Lens Intrinsic membrane 2 protein Lim2 on four subterranean mammal branches. We use maximum likelihood approaches to estimate the amount of sequence divergence of each genetic element on branches of the phylogenetic tree (Yang, 2007). Using the branch lengths on each elements tree's branch lengths, we calculate the average tree across the individual trees reflecting the expected amount of divergence on each branch. Relative evolutionary rates (RERs) on individual trees are then calculated as the residuals of a linear regression analysis where the dependent variable corresponds to the branch lengths of individual trees, and the independent variable corresponds to branch lengths of the average tree. Thus the relative rates reflect the gene-specific rate of divergence in each branch, factoring out the expected divergence on the branch due to genome-wide effects (such as mutation rate, time since speciation, etc.). The relative rates method works downstream of estimating the trees, and hence considers protein-coding gene trees, non-coding genetic element trees and simulated gene trees equivalently. For the sake of simplicity, we refer to the relative rates on the branches of each tree as the gene-specific relative rate; the term gene could in principle be referring to a protein-coding gene, non-coding genetic element, or a simulated tree depending on the dataset being studied.

**Figure 1.**
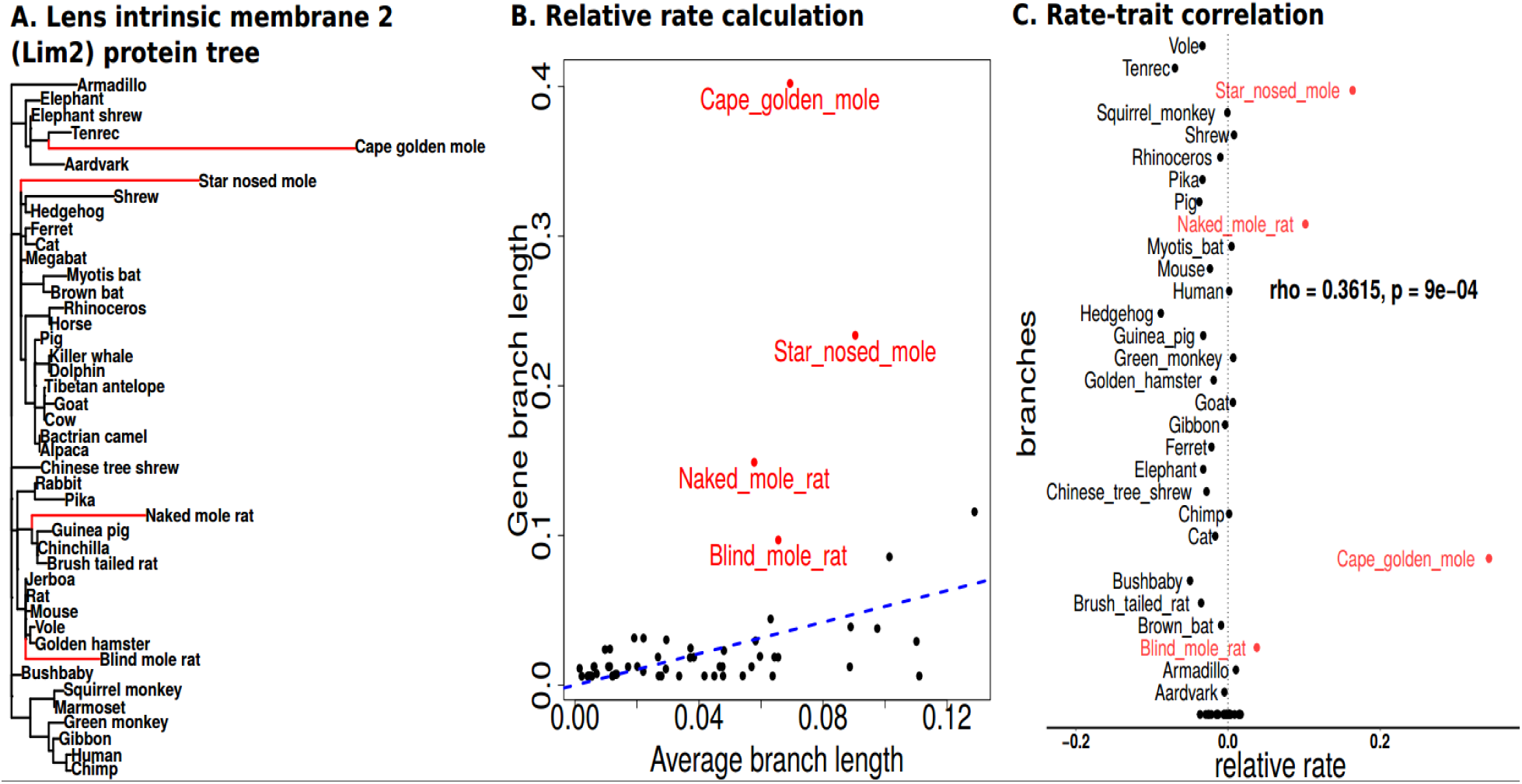
Predicting gene-trait associations using relative rates methods. A. Lens Intrinsic Membrane 2 (Lim2) protein-coding gene tree. Our phylogenetic dataset is comprised of trees constructed from alignments of protein-coding genes in the mammalian genome across 59 species of placental mammals. B. Relative rates on branches of phylogenetic trees are calculated using linear regression. C. Gene-trait associations are identified using correlations of relative rates of the gene with binary trait of interest

### Estimating mean-variance trends in relative rates

Our current methods calculate the gene-specific rates by correcting for the genome-wide effects on branch lengths using linear regression. Consequently, the variance of the relative rates on individual branches strongly depends on the average length of the branch, illustrated here using an example protein-coding gene tree for MFNG, Manic Fringe Homolog Drosophila (Figure 2A). We see that longer branches have relative rates showing a higher variance, as can be inferred from the increasing spread of the relative rates. This pattern becomes clearer when we plot the genome-wide variance in relative rates for branches of different average lengths (Figure 2B). In statistical terms, the relative rates are heteroscedastic meaning they show unequal variance across the range of values of the dependent variable, here the average branch length. The presence of a non-constant mean-variance trend in the residuals stands in violation of one of the assumptions underlying linear regression, namely, homoscedasticity or constant variance of residuals with respect to the dependent variable. More importantly, we suspect that this heteroscedasticity of the relative rates adversely affects the confidence with which we can infer rate shifts on specific branches. For example, the presence of a mean-variance trend can increase the likelihood of observing higher relative rates on longer branches by chance, rather than due to gene-specific changes reflecting changes in selective pressure. A potential negative consequence could be a higher proportion of false positives while inferring convergent rate changes on such branches.

**Figure 2.**
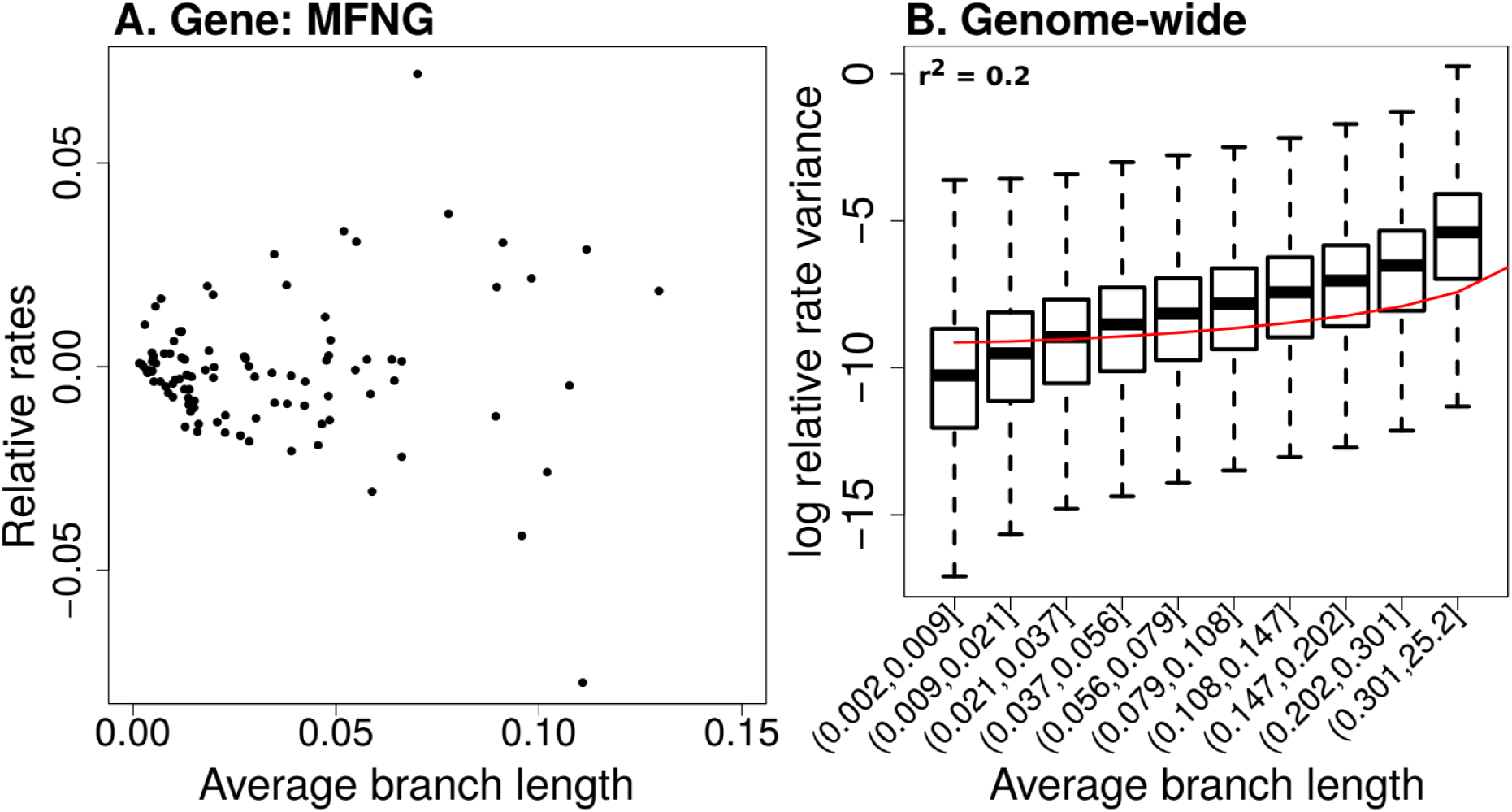
Heteroscedasticity in the relative rates computed using current method. A. Relative rates on branches of Manic Fringe (MFNG) gene tree, calculated using current method. Points represent branches of the gene tree, with relative rates computed on the branches plotted against the genome-wide average length. Heteroscedasticity in the relative rates can be visualized as the increase in the variance of the relative rates with increasing average branch length B. Genome-wide mean-variance trends in relative rates. The logarithm of the relative rate variance within each bin is shown, where branches are binned based on their average lengths across all gene trees. Bin ranges were chosen to provide equal numbers of observations per bin. Higher variance in relative rates are observed with increasing branch lengths, and the extent of this heteroscedasticity is calculated using the ‘r-squared’ of the quadratic model between the variables plotted.

### Updated method to calculate relative rates

In this study, we present an approach relying on a combination of data transformation and weighted linear regression to calculate relative evolutionary rates that addresses the statistical limitations resulting from relative rates calculated using naive linear regression. The proposed method updates are based on the ideas presented in Law et al, who developed new linear modeling strategies to handle issues related to mean-variance relationship of log-counts in RNA-seq reads (Law *et al.*, 2014; Ritchie *et al.*, 2015). We represent the branch lengths on individual gene trees as a matrix Y, where rows correspond to individual genes (g), and columns to the branches (b) on these trees. We first transform the branch length data using a square-root transformation (Eq. 1).

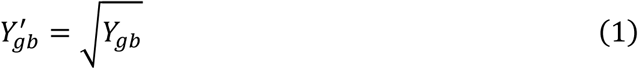

Following the transformation, we perform a weighted regression analysis to calculate the relative evolutionary rates as follows: we calculate the average tree and perform a first-pass of linear regression using the transformed branch length matrix (Eq. 3,4).

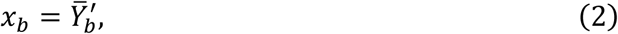

where *x_b_* is the branch length for branch b in the average tree.

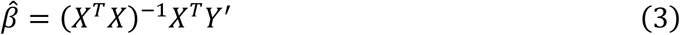

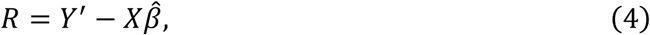

where 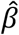 are the coefficients of linear regression, and R is the residuals matrix.

We then estimate the mean-variance trends in the residuals of the linear regression analysis by empirically fitting a locally weighted scatterplot smoothing (LOWESS) function capturing the relationship between the log of variance of the residuals and the branch lengths (Eq. 5).

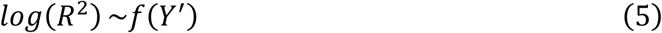

Subsequent to estimating this function, we assign each gene x branch observation a weight W based on the predicted value for the branch, obtained from the first pass linear regression (Eq. 6).

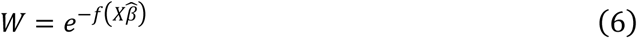

For branches that are shorter on average, the variance in the residuals is smaller, thus resulting in a higher weight, and vice versa. Using the computed weights, we perform a weighted regression analysis between the individual branch length (dependent variable) and the average tree (independent variable). The weighted regression analysis attempts to remove the heteroscedasticity in the residuals by computing the residuals after minimizing the weighted sum of squared errors, as opposed to the raw sum of squared errors (Eq. 7,8).

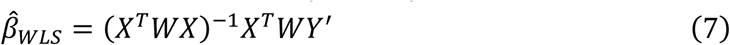

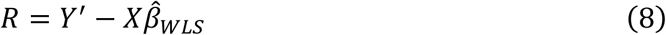

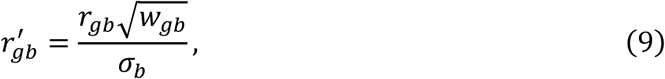

where *σ_b_* is the standard deviation of the weighted residuals in branch b

Subsequent to the weighted regression analysis, the weighted residuals 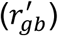, are estimated by rescaling the regression residuals (r_gb_) with the weights, additionally standardized to have unit variance within every branch across all genes (Eq. 9). The weighted residuals 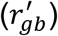 correspond to the weighted relative rate on branch *b* for gene *g*. The differences to the relative rate calculations introduced by the updated method result in changes to the scales of the relative rates computed. However, we note that this scale is arbitrary and the downstream gene-trait correlations for binary traits estimated using a Mann-Whitney test (see Methods) depend only on the ranks of the relative rates of each branch within any single gene tree. Figure 3 shows the workflows for computing relative evolutionary rates using the original and updated method.

**Figure 3.**
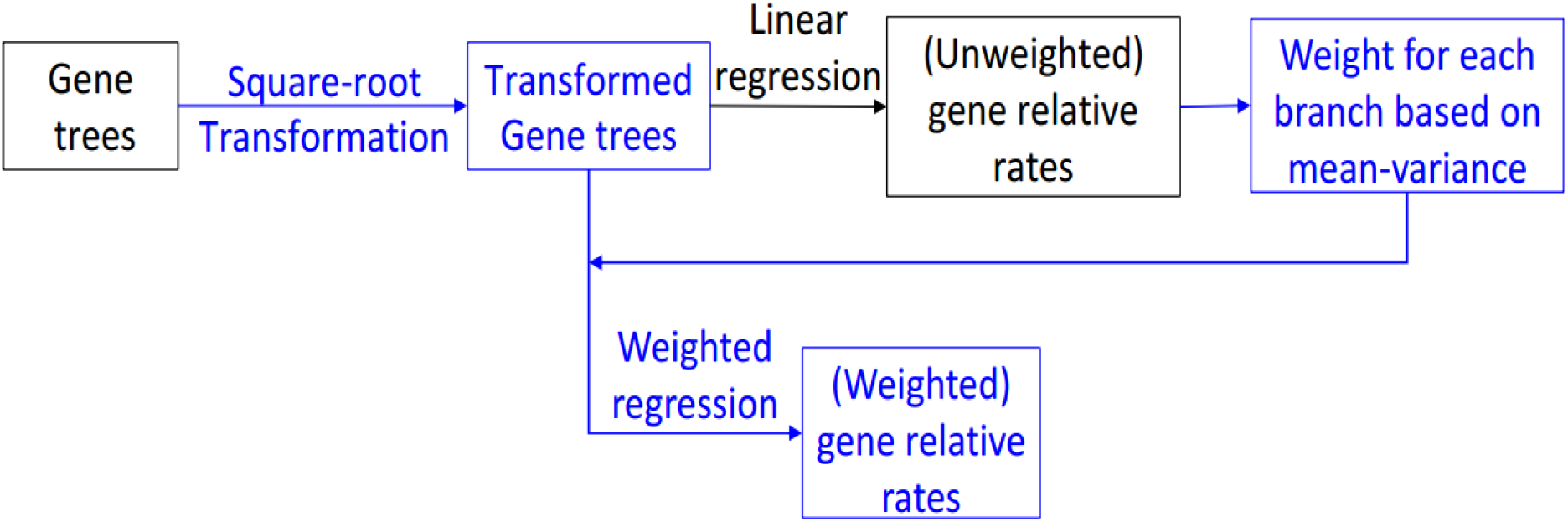
Workflow for calculating relative evolutionary rates using the updated method. Black areas of the workflow represent steps implemented as part of current relative rates method, and blue areas correspond to methodological updates.

## Results

### Improvements to relative evolutionary rates methods mitigate genome-wide mean-variance relationship

Our updated method to calculate relative rates using data transformation followed by weighted regression produce nearly homoscedastic relative rates that do not show a significant global mean-variance relationship. Figure 4A shows the relative rates computed for the MFNG protein-coding gene tree using the updated method. In comparison to the original method based on naive linear regression (Figure 2A), we observe that the updated method produces relative rates showing no apparent increase in the variance of relative rates on longer branches of the tree. Plotting the genome-wide mean-variance trends of the relative rates, we observe that the relative rates calculated from transformed-weighted residuals show nearly constant variance across branches of varying lengths (Figure 4B). We additionally checked the mean-variance relationships from intermediate steps in our method that can estimate relative rates, corresponding to two method variants which do not implement data transformation (linear-weighted regime) or a weighted regression (square-root unweighted regime) (Supplementary Figure S1). However, we find that the intermediate regimes, utilizing only one of the method updates (branch length transformation or weighted regression) are less effective at eliminating mean-variance trends. A combination of transformation and weighted regression steps works best at producing homoscedastic relative rates.

**Figure 4.**
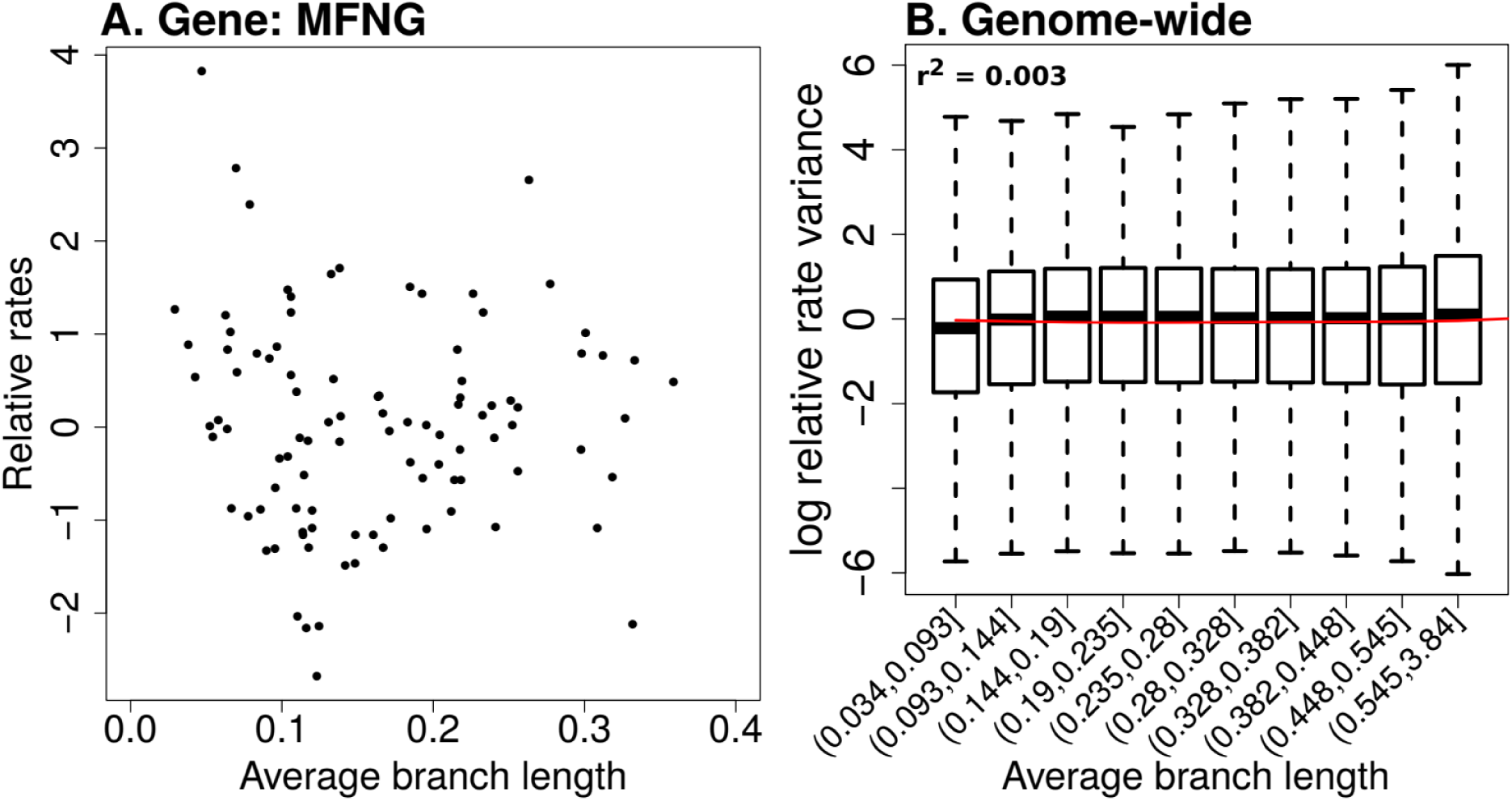
Updated method to calculate relative rates shows no apparent trends of heteroscedasticity. A. Manic Fringe (MFNG) gene relative rates calculated using the updated method. In comparison to figure 2A, we do not observe an increase in the variance of relative rates of branches with increasing average branch length. B. Genome-wide mean-variance trends for relative rates computed using the updated method show constant variance with increasing branch lengths. Contrasting the trends resulting from the application of original (Figure 2B) and updated method (Figure 4B), we observe that the updated method produces nearly homoscedastic relative rates. The extent of heteroscedasticity, computed as the ‘r-squared’ of the quadratic model between the variables plotted, is nearly hundred-fold lower with the updated method compared to original method.

### Better robustness to inclusion of distantly related species

In earlier applications of our evolutionary-rates-based method to detect genetic elements convergently responding in subterranean mammals and marine mammals respectively, we sampled alignments of placental mammal species to construct phylogenetic trees for each genetic element (Chikina, Robinson and Clark, 2016; Partha *et al.*, 2017). These alignments were derived from the placental mammal subset of the 100-way vertebrate alignments made publicly available by the UCSC genome browser (Casper *et al.*, 2018). In addition to these placental mammals, the 100-way alignments include four other species of mammals, three marsupials – Opossum (*monDom5*), Wallaby (*macEug2*), Tasmanian Devil (*sarHar1*), and one monotreme – Platypus (*ornAna1).* Despite deep conservation of many genetic elements in these non-placental mammals, human-and-mouse centered phylogenomic studies tend to exclude these species due to the introduction of long branches in the phylogenetic trees (Parker *et al.*, 2013; Marcovitz, Jia and Bejerano, 2016; Prudent *et al.*, 2016). For instance, in previous applications of our relative-rates-methods we deliberately excluded these non-placental mammals since they produce wide variations in relative rates due to the introduction of long branches, which would adversely affect the confidence with which we make inferences of convergent rate acceleration in species exhibiting a convergent phenotype (Chikina, Robinson and Clark, 2016; Partha *et al.*, 2017). However, scanning for rate-trait associations across tree datasets with higher numbers of species would allow for more statistical power, and hence a relative rates method that can reliably include such distantly related species offers a clear advantage. To this end, we test the robustness of our updated method to the inclusion of distantly related species at inferring convergent rate shifts. We choose two phylogenetic datasets - 1. Genome-wide protein-coding gene alignments across 59 placental mammal species, and 2. across 63 mammals including the four non-placental mammals in addition to the placentals. An example demonstration of how our current method to calculate relative rates is sensitive to the inclusion of non-placental mammals is illustrated in Figure 5A. Using the example *Peropsin* (RRH) gene, we show that the ranks of relative rates computed using the current method considerable vary upon the inclusion of non-placental mammals. These changes in ranks are observed across many branches on the gene tree including one of the four subterranean branches (Cape golden mole). In comparison, the updated method displays a stronger concordance in the ranks of the computed relative rates (Figure 5A). Consequently, the subterranean acceleration scores for RRH computed using the updated method are more stable with the inclusion of non-placental mammals (Table 1).

**Table 1.** Subterranean acceleration scores for Peropsin (RRH) computed using two methods, and across two datasets. In comparison to the original method, the updated method shows stronger consistency in the scores across the two tree datasets, with and without the non-placental mammals. The subterranean acceleration scores reflect the significance of convergent rate acceleration on the four subterranean branches.

| Dataset\Method | Original | Updated |
| --- | --- | --- |
| With non-placentals | 2.70(rho = 0.31;<br>p = 0.002) | 2.1(rho = 0.27;<br>p = 0.008) |
| Placentals only | 1.38(rho = 0.21;<br>p = 0.041) | 2.0(rho = 0.26;<br>p = 0.01) |

**Figure 5.**
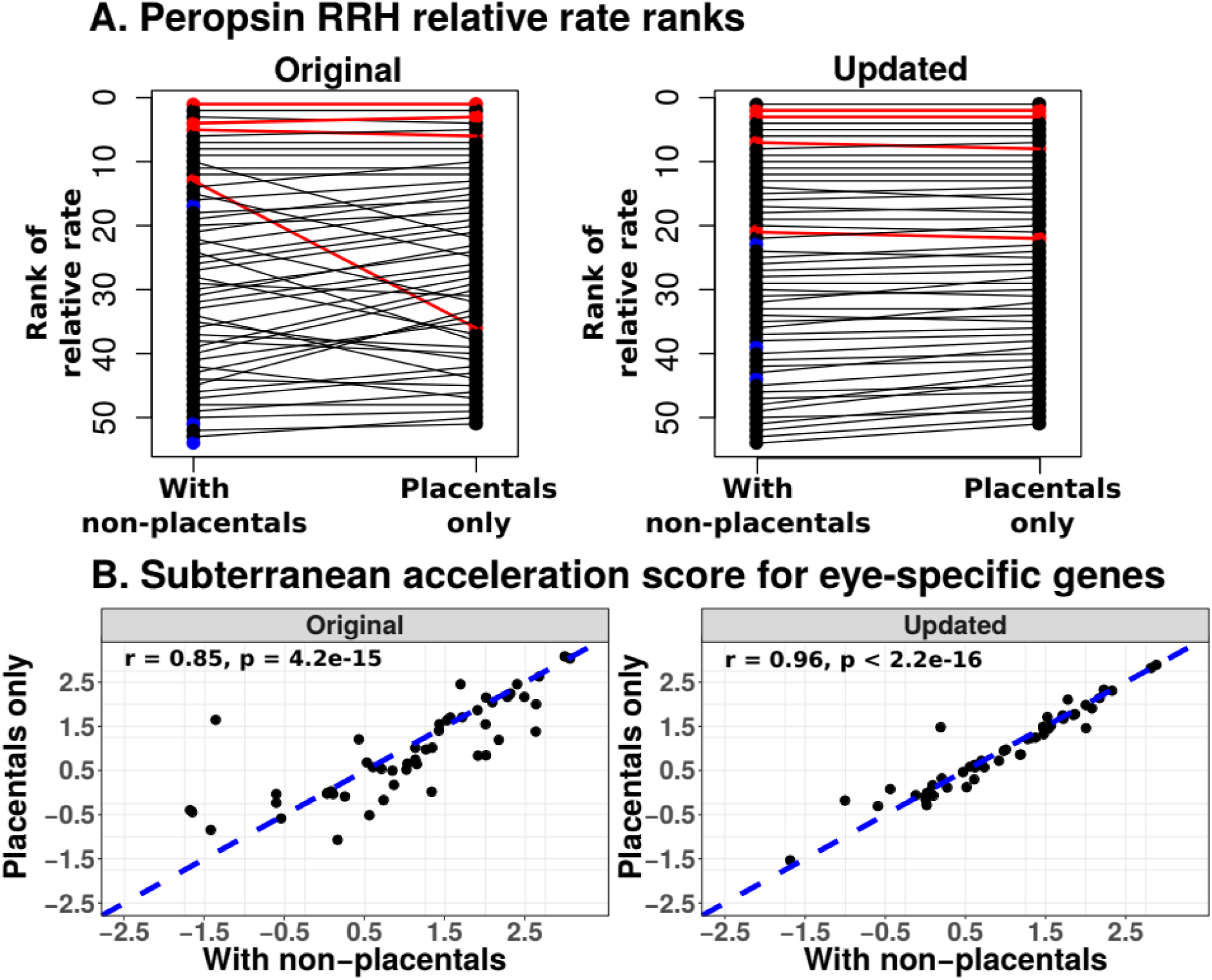
Comparison of robustness of methods to inclusion of non-placental mammals. A. Relative rates of Peropsin (RRH) gene across trees with and without non-placental mammals, using the original vs the updated method. The relative rate ranks of terminal lineage branches within the RRH tree are plotted with respect to the inclusion of non-placental mammals in the gene trees. Red points denote subterranean branches, and blue points correspond to non-placental mammals. Ranks of relative rates on computed by the original method form a crisscross pattern reflecting wider variation with respect to the inclusion of non-placental mammals, whereas updated methods reveal stronger concordance. B. Updated method shows improved robustness to inclusion of non-placental mammals at detecting subterranean acceleration of eye-specific genes. Individual points represent rate acceleration on subterranean branches for each of sixty eye-specific genes computed across two datasets using the two methods. Robustness of each method is estimated as the spearman correlation coefficient of subterranean acceleration scores computed based on placental-only gene trees (y-axis) and scores computed based on trees also including non-placental mammals. An improved robustness to the inclusion of non-placental mammals is observed with the updated method.

We also performed a larger-scale benchmarking of the robustness of our methods to the inclusion of non-placental mammals across 60 genes showing eye-specific expression. These genes were identified based on mouse microarray expression data across 91 tissues (see Methods). For each of these eye-specific genes, we calculated subterranean acceleration scores (see Methods) reflecting the convergent rate acceleration on the four subterranean branches, independently in gene trees including and excluding the non-placental mammals. Based on the relative rates calculated using each method, we compared the concordance of the subterranean acceleration scores across the two tree datasets. Ideally, we expect the scores produced by the methods to be highly consistent across the two datasets since the four non-placental mammals are not subterranean by nature, with only minor differences arising due to the inclusion of four additional background species. The results of the analysis revealed that the updated method produces superior concordance in the scores across the two tree datasets, reflecting its improved ability to handle the long branches introduced by the non-placental mammals, (Fig 5B).

### Improved power to detect convergent rate shifts in simulated trees

In order to compare the power of our methods to detect convergent rate shifts in branches across a range of evolutionary scenarios, we developed a model to simulate individual gene trees. Such a model allows us to rigorously examine method performance in relation to various parameters in phylogenetic datasets including number of foreground branches, length distribution of foreground branches etc., where foreground branches describe branches showing a convergent phenotype, while background branches do not. The limited availability of ‘ground truth’ examples of convergently evolving genetic elements calls for the development of biologically realistic simulations of sequence evolution. Using our model to simulate trees (see Methods), we compared the power to detect rate shifts in relation to two factors: 1. Average lengths of foreground branches, in particular extreme foreground branches - branches that are short and long on average, respectively. 2. Number of foreground branches. We investigated the performance of the updated method in detecting rate shifts in such extreme branches, assessing the power advantage resulting from calculating relative rates that do not suffer from a biased mean-variance relationship.

Our model to simulate phylogenetic trees allows for explicit control over choosing foreground branches showing convergent rate acceleration. We simulate ‘control’ trees, where all branches are modeled to evolve at their respective average rates, and ‘positive’ trees, where the chosen foreground branches are modeled to evolve at an accelerated rate that is twice their average rate (see Methods). Firstly, we compared the heteroscedasticity in the relative rates on the branches of the control trees calculated using the original and updated methods. Similar to the trends observed in mammalian gene trees (Supplementary figure S1), we observed that the updated method outperformed the original method at producing homoscedastic relative rates (Supplementary figure S3). We then calculated a foreground acceleration score for individual simulated trees (see Methods). This score, calculated a signed negative logarithm of the p-value, is higher for trees showing stronger convergent rate acceleration on the foreground branches. Subsequent to estimating these scores, we evaluated the performance of the two methods, based on the power to distinguish the positive trees as showing convergent acceleration in comparison to the control trees. In two independent simulation settings with foreground branches of long and short average lengths respectively, we observed that the updated method offers more power to detect convergently accelerated ‘positive’ trees (Figure 6B). In addition to the positive trees with foreground branches that were long or short branches, we compared the power to detect rate acceleration on foreground branches of intermediate length. Consistent with the findings in short/long foregrounds, we find a modest yet significant improvement offered by the updated method (Supplementary figure S4). Overall, we find that our updated method to compute relative rates offers a significantly improved power to detect convergent rate shifts in simulated trees.

**Figure 6.**
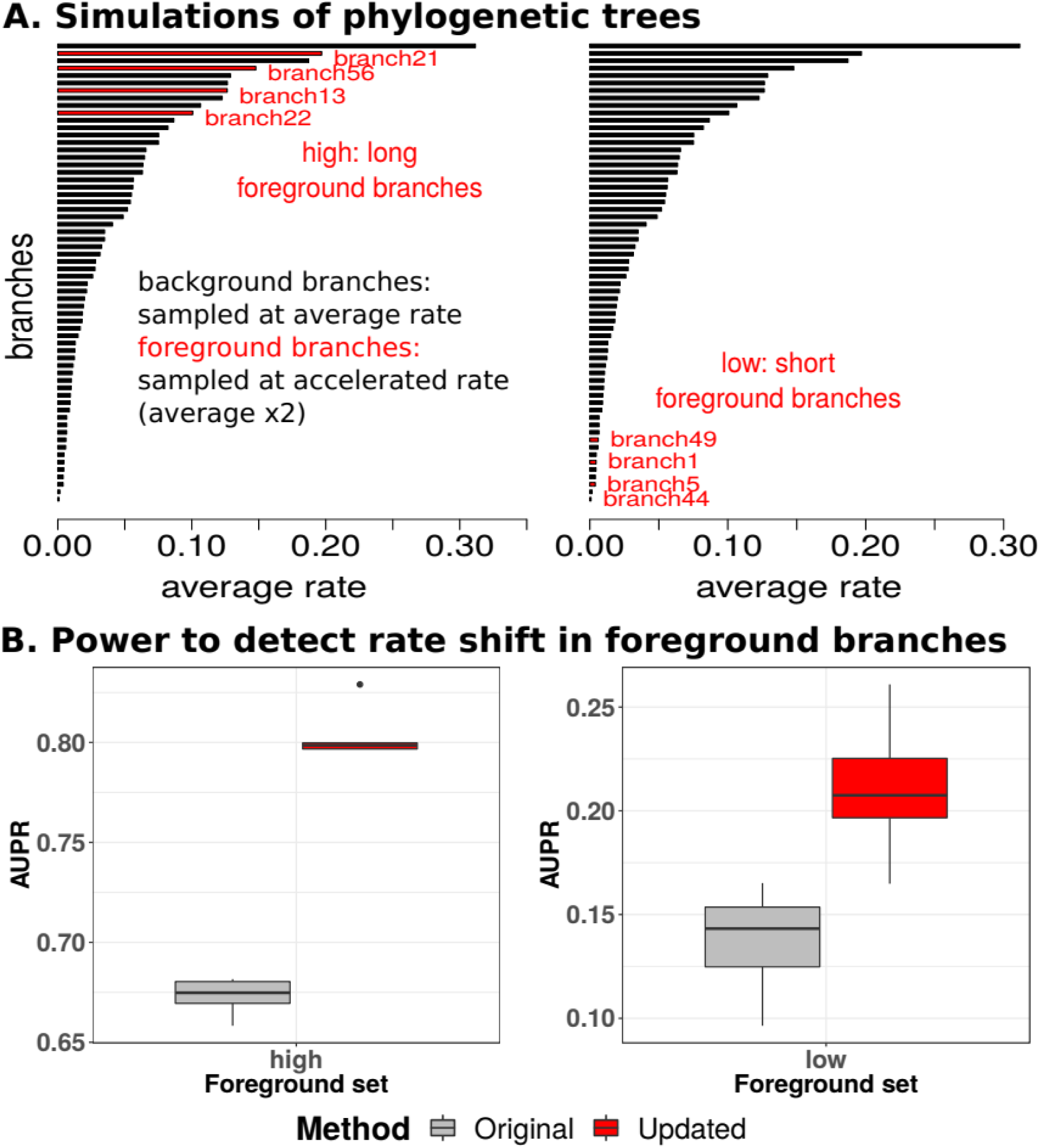
Comparison of method performance across simulated phylogenetic trees. A. Branch length distributions for simulating phylogenetic trees with foreground branches highlighted in red. Two independent simulations were performed with foreground branch sets comprised of long foreground branches (left panel) and short foreground branches (right panel) respectively. B. Power to detect rate shift in foreground branches of simulated trees. Across five independent simulations of control trees and positive trees, we measured the area under the precision-recall curve (AUPR) to precisely detect positive trees using the foreground acceleration score. The AUPR distributions obtained using the updated method to calculate relative rates are significantly elevated compared to the original method across simulated scenarios involving foreground sets of long (left) and short branches (right) respectively.

We then compared the power to detect rate shifts across varying numbers of foreground branches by simulating ‘positive’ trees with seven foreground branches of short average lengths (Supplementary figure S5). We subsequently generated positive trees with subsets of n branches (n ranging from 4 to 7) among these seven foreground branches (Supplementary figure S5). Within each of these datasets, we calculated foreground acceleration scores for control and positive trees using each method independently. Contrasting the power of the methods to precisely detect positive trees as showing convergent rate acceleration, we observed that the updated method to calculate relative rates is consistently more powerful than the original method. We repeated the analysis choosing seven foreground branches that were long on average rather than short (Supplementary figure S5). Similar to the short foreground scenario, we observed that the updated method is able to more precisely detect convergent rate acceleration with the inclusion of additional foreground branches in the simulated trees (Figure 7B).

**Figure 7.**
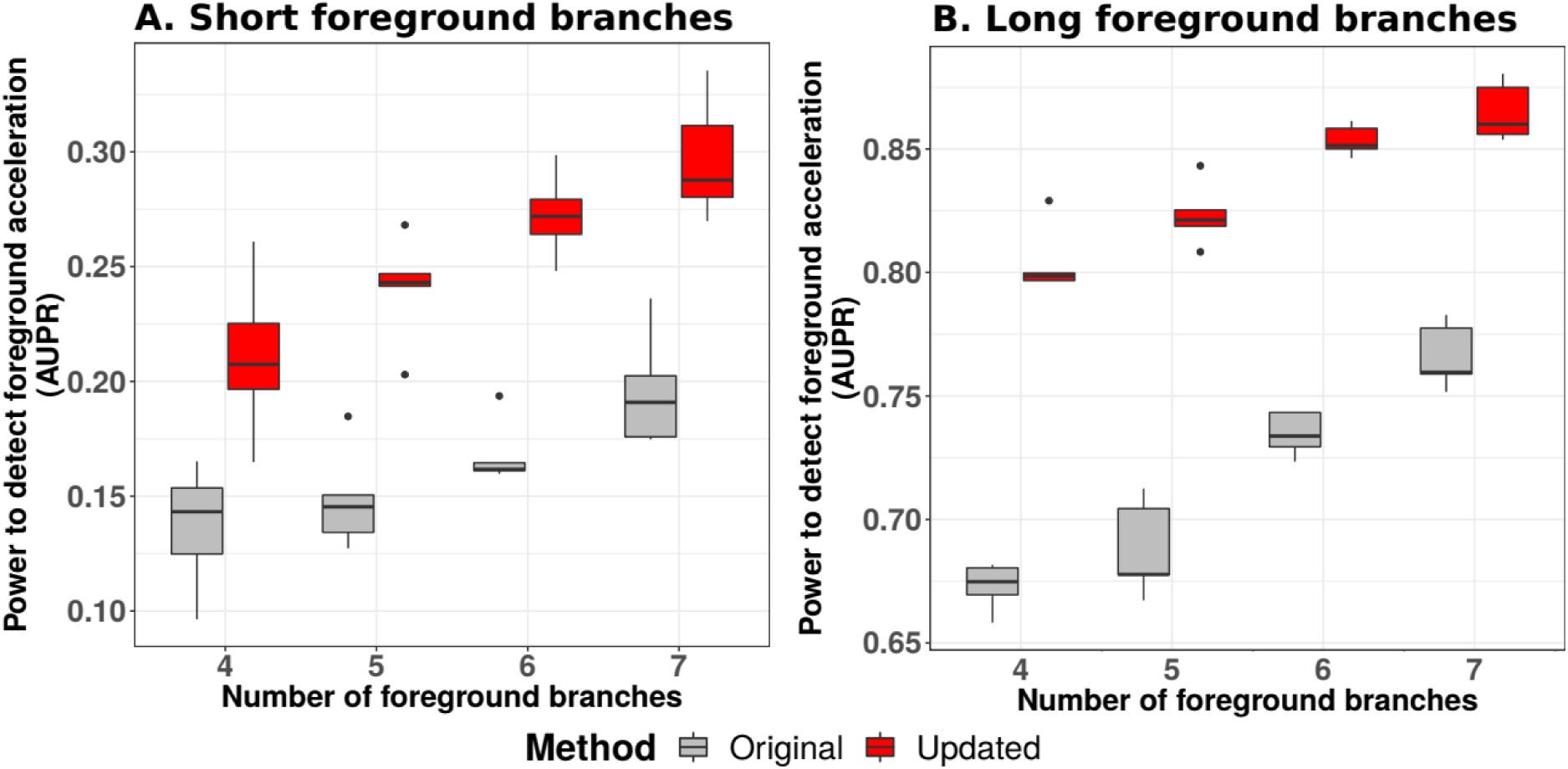
Improved power to detect foreground rate shifts using the updated method across different numbers of foreground branches. This analysis was performed across five independent simulations of control trees and positive trees with varying numbers of foreground branches (4 to 7). Within each simulation, we measured the area under the precision-recall curve (AUPR) to precisely detect positive trees using the foreground acceleration score. The AUPR distributions obtained using the updated method to calculate relative rates are consistently elevated compared to the original method across simulations with different numbers of foreground branches. These simulations were performed across two scenarios with different foreground branch sets consisting short (A) and long branches (B) respectively.

### Relative rates-based inference is robust to minor uncertainties in species tree topology

Our method relies on estimating sequence divergence on branches of phylogenetic trees with a fixed topology. Efforts to better resolve the phylogeny of extant mammals have resulted in continuous updates to the consensus species tree topology (Murphy *et al.*, 2001, 2007). Topology trees commonly used in phylogenomic analyses of extant mammals include the UCSC genome browser’s 100-way tree, as well as the timetrees reported in the Meredith et al and Bininda-Emonds et al (Bininda-Emonds *et al.*, 2007; Meredith *et al.*, 2011; Casper *et al.*, 2018). Differences between these species tree topologies often involve entire clades, and the decision to choose a particular topology tree can potentially strongly influence the outcomes of phylogenetic analyses. Here, we benchmarked the robustness of our relative rates methods to the choice of topology tree. We constructed protein-coding gene trees based on two different species tree topologies, namely the UCSC 100-way tree and our modified Meredith et al. (Meredith+) topology tree (see Methods). The Robinson-Foulds metric between these two phylogenies is 22, reflecting differences in 22 partitions of species (Robinson and Foulds, 1981; Schliep, 2011). We observed that both the updated and original methods to calculate relative rates show robust signatures of subterranean rate acceleration for eye-specific genes with respect to the species tree topology used (Figure 8).

**Figure 8.**
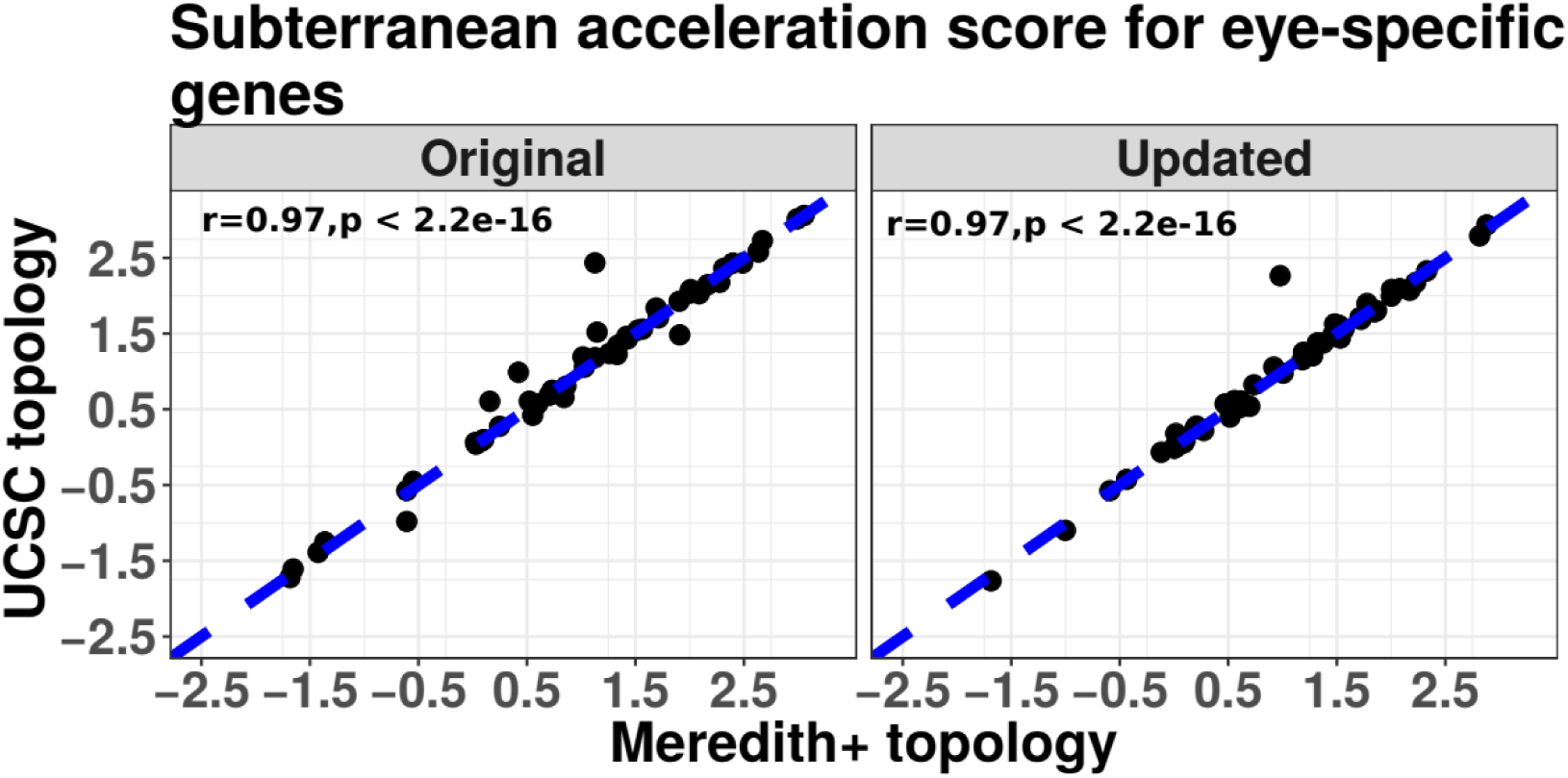
Comparison of robustness of methods to species tree topology. Both the original and updated relative rates based methods are robust to choice of species tree topology used to construct individual gene trees. Points represent the strength of convergent subterranean acceleration for eye-specific genes whose trees were constructed using the Meredith+ topology (x-axis), and the UCSC topology (y-axis) respectively. The robustness of each method is estimated as the spearman correlation coefficient of the subterranean acceleration scores across the two tree datasets.

## Discussion

Our previous evolutionary-rates-based method to detect genomic elements underlying convergent phenotypes has already proved to be a valuable technique to detect genes and enhancers associated with transitions to marine and subterranean habitats (Chikina, Robinson and Clark, 2016; Partha *et al.*, 2017). However, the original method suffered from reduced power to detect such genomic elements due to a heteroscedastic relationship between the mean and variance of branch lengths for a given branch across all gene trees, i.e. branches that are longer on average have higher variance than branches that are shorter on average. Here, we developed a method using a square-root transformation and a weighted regression based on the observed mean-variance relationship to correct for the heteroscedasticity. We tested our improved method on real and simulated phylogenies and observed improved robustness to wider ranges of branch lengths and increased ability to detect convergent evolutionary rate shifts. Our method, called RERconverge, is freely-available on GitHub (Kowalczyk *et al.*, 2018) (https://github.com/nclark-lab/RERconverge).

Our new method offers increased robustness to the inclusion of distantly-related species with long branch lengths in our phylogeny, namely non-placental mammals. When we compared results from an analysis using only placental mammals and an analysis that included non-placental mammals using both our previous and our updated methods, we found that our new method, unlike the previous method, is unimpaired by the inclusion of non-placental mammals. By improving our method’s robustness to inclusion of long branches, we increased the method’s applicability to a broader range of species and hence a broader range of convergent phenotypes. Additionally, our new method’s increased power could enable us to discover more convergently evolving genomic elements. One particular incentivizing example for these improvements is the recent efforts to sequence the northern marsupial mole, a completely blind mammal (Archer *et al.*, 2011). When considering using subterranean species to find genes and enhancers associated with vision, the ability to include the non-placental marsupial mole along with the other non-placental mammals in our dataset will allow for more power in a scan for vision-specific genetic elements showing convergent regression in the five blind mammals.

Beyond examining individual genes, we further assessed our new method’s ability to detect pathway enrichments for genes under relaxation of constraint in subterranean mammals and marine mammals (see Figure 9 for marine foreground branches). Compared to our previous method, the updated method detected more enriched Gene Ontology (GO) terms with accelerated evolutionary rates in subterranean mammals (Table 2). Additionally, the fold enrichment for detected terms was significantly stronger with the updated method (Figure S6, Supplementary Table S1). On the other hand, the marine system showed mixed results. Both the new and the old method showed approximately equal power to detect enriched GO terms if we only consider the number of terms detected (Table 3). However, when comparing the fold enrichment for detected terms, the old method was significantly better than the updated method (Figure S6, Supplementary Table S1). These contrasting results from the subterranean dataset versus the marine dataset indicate the importance of tailoring the corrections we have developed to the dataset of interest, as well as the importance of taking advantage of simulation-based power and robustness assessments to develop methods that are broadly applicable to many convergent phenotypes.

**Table 2.**
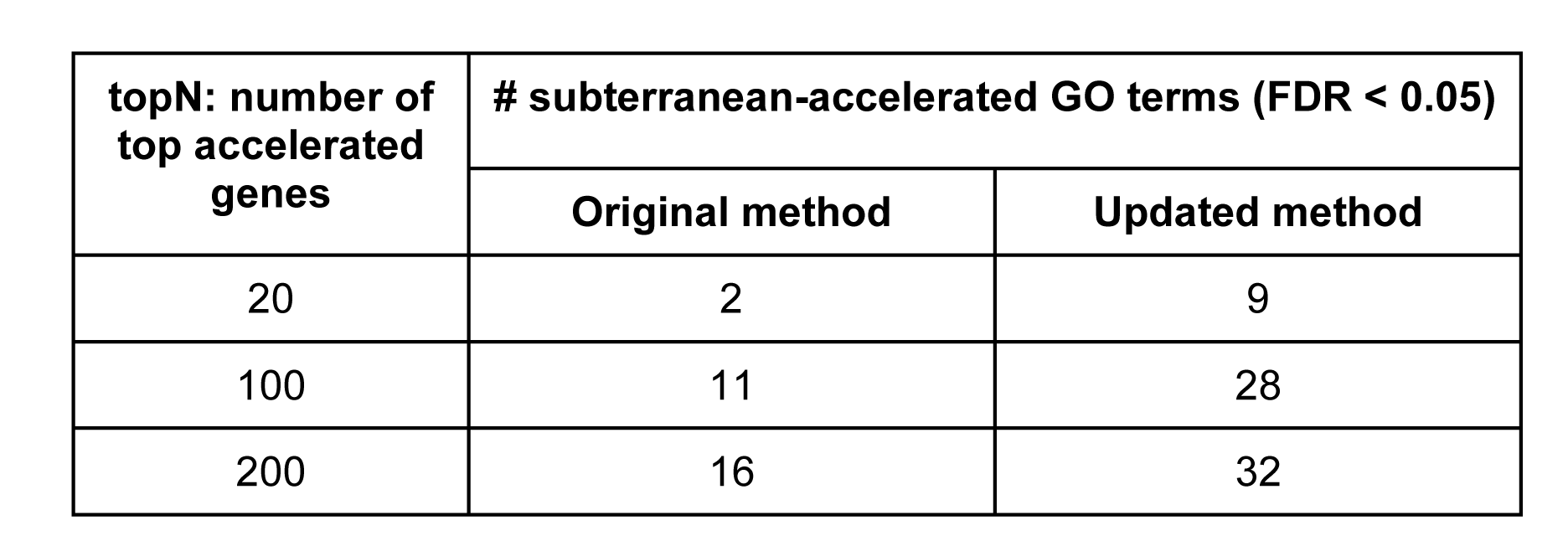
Comparison of number of vision-related Gene Ontology terms enriched in top subterranean-accelerated genes discovered by the original and updated methods. Gene Ontology term enrichment analysis was performed individually on top subterranean accelerated genes discovered by each method. Across varying numbers of top target genes, genes discovered using the updated method were consistently enriched for higher numbers of vision-related GO terms.

| topN: number of top accelerated genes | # subterranean-accelerated GO terms (FDR < 0.05) |  |
| --- | --- | --- |
|  | Original method | Updated method |
| 20 | 2 | 9 |
| 100 | 11 | 28 |
| 200 | 16 | 32 |

**Table 3.** Comparison of number of Gene Ontology terms enriched in top marine-accelerated genes discovered by the original and updated methods. Gene Ontology term enrichment analysis was performed individually on top marine-accelerated genes discovered by each method. Across varying numbers of top target genes, neither method showed a clear superiority over the other at detecting higher numbers of enriched terms.

| topN: number of top accelerated genes | # marine-accelerated GO terms (FDR < 0.05) |  |
| --- | --- | --- |
|  | Original method | Updated method |
| 50 | 16 | 10 |
| 100 | 27 | 31 |
| 200 | 59 | 59 |

**Figure 9.**
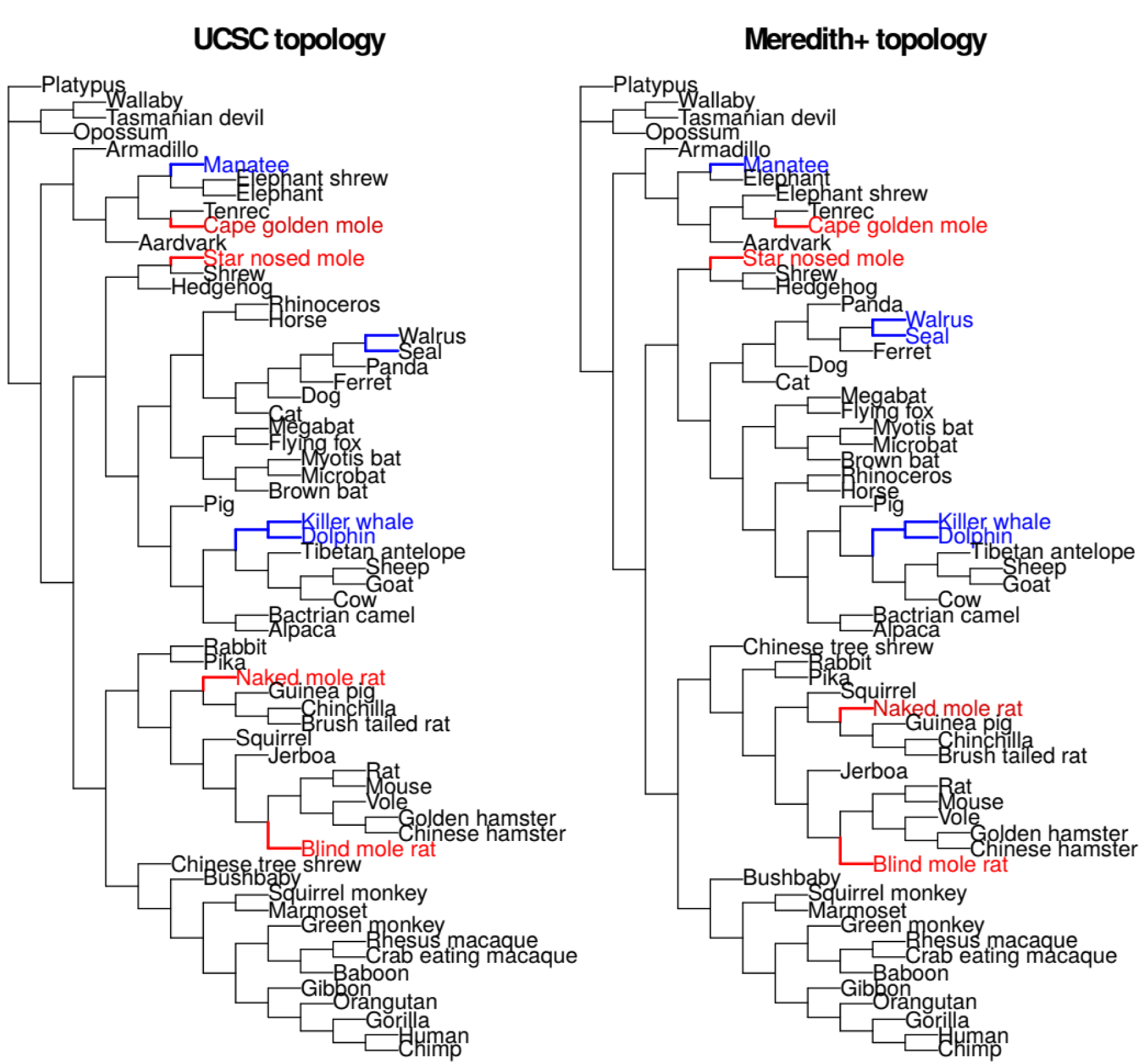
Cladograms describing relationships between 63 mammalian species used for constructing genome-wide maximum likelihood protein-coding gene trees. Tree reported in UCSC genome browser (left) and the final version of the tree we modified from the topology reported in Meredith et al (right). Branches corresponding to subterranean and marine mammals are highlighted in red and blue, respectively.

In addition to testing our method on real data, we also developed a simulation-based strategy to represent a "true positive” case of convergent evolution. Our simulations follow a similar approach to simulating RNA-seq counts where simulated rates are essentially capturing the number of substitutions that occur along a branch (Di *et al.*, 2011). We showed that our new method demonstrates improved detection of rate shifts both when foreground species occupy long, high-variance branches and when foreground species occupy short, low-variance branches. This allows the method to detect convergent rate shifts given a variety of potential configurations of convergently-evolving species. The types of simulations we developed are essential because relatively few concrete instances of sequence-level evolutionary convergence exist, so biologically accurate simulations of such evolution are essential to rigorously test methods that detect shifts in evolutionary rates. One simplification of our simulation method is that all species are present in all simulated trees, which is not the case in real genomic data because of genomic element gain and loss across species. However, maintaining constant species composition in our simulated trees should have little impact on our ability to compare our methods because we expect both to be equally impacted by species presence and absence. A second simplification is that we assume all convergently-evolving species have the same phylogenetic relatedness, i.e. each foreground branch is an independent instance of convergent evolutionary rates. We would like to be able to answer questions about our method’s power given more complex phylogenetic configurations. Developing methods to answer those types of questions will require a much higher degree of complexity in our simulations, but it will also allow us to determine which species to add to our genomic datasets to increase our power to find convergently evolving genomic features.

Our improved method has proved valuable for detecting genomic elements associated with two binary traits - subterranean-dwelling or not, and marine-dwelling or not - and we will extend our method for use in convergent continuous traits and non-binary discrete traits. We will also assemble complementary analyses to assess the robustness and power of each method. By extending the scope of our method to non-binary traits, we will expand the potential search-space of our method to a plethora of new convergent phenotypes. Our overarching goal is to develop an entire suite of methods that can utilize any conceivable phenotypes as inputs to accurately and robustly identify convergently evolving genomic elements.

## Materials and Methods

### Protein-coding gene trees across 63 mammalian species

We downloaded the 100-species multiz amino acid alignments available at the UCSC genome browser, and retained only alignments with a minimum of 10 species. We then pruned each alignment down to the species represented in Figure 9 of the proteome-wide average tree. We added the blind mole rat ortholog of each gene based on the methods described in Partha et al (Partha *et al.*, 2017). We estimated the branch lengths for each amino acid alignment using the *aaml* program from the package PAML (Yang, 2007). We estimated these branch lengths on a tree topology modified from the timetree published in Meredith et al. We attempted to resolve conflicts between the topology inferred in Meredith et al. compared to that in Bininda-Emonds et al. based on a consensus of various studies employing a finer scale phylogenetic inference of the species involved (Bininda-Emonds *et al.*, 2007; Meredith *et al.*, 2011). The differences between our final topology, which we call ‘Meredith+’ topology and the Meredith et al. topology include setting the star-nosed mole as an outgroup to the hedgehog and shrew; cow as an outgroup to the Tibetan antelope, sheep and goat; and the ursid clade as an outgroup to mustelid and pinniped clades. For more details about the literature surveyed to resolve these differences, please refer to Meyer et al (Meyer *et al.*, 2018). The topology of our final ‘Meredith+’ tree compared to the UCSC topology tree is reported in Figure 9. In order to perform analyses benchmarking the method robustness to tree topology, we additionally generated the protein-coding gene trees based on the UCSC tree topology.

### Simulating phylogenetic trees

Phylogenetic branch lengths have units of number of substitutions per site and thus can be thought of as normalized count data. However, we find that a *Poisson* distribution is unsuitable in this case as the real branch length data shows considerable overdispersion, that is the variance is higher than the mean. We thus model the branch lengths of the simulated trees using a negative binomial distribution, following ideas from studies simulating expression counts for RNAseq analysis (Robinson, McCarthy and Smyth, 2009; Di *et al.*, 2011; Law *et al.*, 2014; Ritchie *et al.*, 2015).

We simulated datasets of phylogenetic trees using the UCSC tree topology and branch lengths from the average proteome-wide tree across 19,149 mammalian protein-coding gene trees across 62 mammals. Supplementary figure S2 describes the tree topology used for the simulations. We simulate the branch lengths (or rates) for every branch (*j*) on each tree (*i*) according to the following formula,

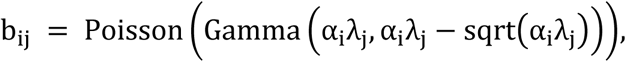

where *Gamma* is parametrized by mean and variance. Here, *α_i_* is a gene-specific scaling term, *λ_j_* is the average rate of the corresponding branch so that *α_i_λ_j_* is the expected rate on the *ij*’th branch, and the simulated rate is drawn from a *Gamma* distribution with that mean. The composite *Poisson-Gamma* distribution is equivalent to the negative binomial distribution and thus in our simulation the mean variance relationship has a quadratic component, matching what we observe in real data.

We simulate two classes of trees in every dataset based on different input parameters. We simulate ‘control’ trees, trees where the j are simply the average rate on the branch j. These ‘control’ trees do not show any explicit convergent rate shift on any of the branches. We additionally simulate ‘positive’ trees showing convergent rate acceleration on foreground branches by sampling at 2*λj, only on these branches. Thus, the foreground branches in positive trees are effectively sampled at an accelerated rate compared to the foreground branches in control trees.

### Calculating gene-trait correlations

The gene-trait correlations are computed under a Mann-Whitney U testing framework over the binary variable of foreground vs background branches. In the subterranean example, the four subterranean branches (Figure 1) are designated as foreground. We calculate a foreground acceleration score reflecting the strength of convergent rate acceleration on the foreground branches. The value is calculated as the negative logarithm of the p-value of the Mann-Whitney test multiplied by the direction of the correlation as given by the sign of the rho statistic. A positive rho statistic indicates rate acceleration in the foreground species, and the negative logarithm of p-value reflects the strength of the convergent rate shift. In simulated trees study, we generated trees for three sets of foreground branches with different branch length distributions - short, intermediate, and long as illustrated in Figure 6 and Figure S5.

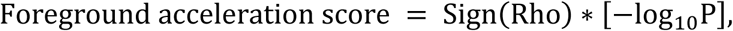

where rho and P are the correlation coefficient and statistical significance of the Mann-Whitney test for association between relative rates and binary trait.

### Genes showing eye-specific expression

We identified eye-specific gene sets using microarray expression data from 91 mouse tissues (Su *et al.*, 2004). We identified genes specifically expressed in the following tissues of the eye - cornea, iris, lens and retina (including retinal pigmented epithelium). These genes showed significant differential expression only in the tissue of interest compared to the other tissues at an alpha of 0.05 (T-test)

### Gene Ontology term enrichment analysis

We performed functional enrichment analysis in target gene lists using the GOrilla tool (Eden *et al.*, 2009). For each analysis, GO terms enriched in target gene lists were identified by comparing to a background gene list with all 19,149 genes used to construct gene trees.

## Acknowledgements

We thank Dennis Kostka, as well as members of the Chikina, and Clark labs for helpful discussion and feedback on earlier versions of this manuscript. This work was supported by NIH R01 HG009299-01A1 to N.L.C and M.C

